# Glucocorticoids Mediate Transcriptome-wide Alternative Polyadenylation: Mechanisms and Functional Implications

**DOI:** 10.1101/2022.03.30.486468

**Authors:** Thanh Thanh Le Nguyen, Duan Liu, Huanyao Gao, Zhenqing Ye, Jeong-Heon Lee, Lixuan Wei, Jia Yu, Lingxin Zhang, Liewei Wang, Tamas Ordog, Richard M. Weinshilboum

## Abstract

Alternative polyadenylation (APA) is a common genetic regulatory mechanism that generates distinct 3′ ends for RNA transcripts. Changes in APA have been associated with multiple biological processes and disease phenotypes. However, the role of hormones and their drug analogs in APA remains largely unknown. In this study, we investigated transcriptome-wide the impact of glucocorticoids on APA in 30 human B-lymphoblastoid cell lines. We found that glucocorticoids could regulate APA of a subset of genes, possibly by changing the expression of 142 RNA-binding proteins, some with known APA-regulating properties. Interestingly, genes with glucocorticoid-mediated APA were enriched in viral translation-related pathways, while genes with glucocorticoid-mediated expression were enriched in interferon and interleukin pathways, suggesting that glucocorticoid-mediated APA might result in functional consequences distinct from gene expression. Glucocorticoid-mediated APA was also cell-type-specific, suggesting an action of glucocorticoids that may be unique to immune regulation. We also observed evidence for genotype-dependent glucocorticoid-mediated APA, providing potential functional mechanisms for a series of common genetic variants that had previously been associated with immune disorders, but without a clear mechanism. In summary, this study reports a series of novel observations regarding the impact of glucocorticoids on APA, suggesting novel avenues for mechanistic studies of hormonal biology.

## INTRODUCTION

Alternative polyadenylation (APA) is a molecular regulatory mechanism that generates RNA isoforms with differential usage of polyadenylation sites (PASs) (1). When APA takes place in the final exon of a gene, it creates RNA transcripts with differential lengths of the 3’-untranslated region (3’-UTR) of the gene. When APA takes place in introns, it can generates transcript variants with new 3’ ends, a process that often involves differential splicing (1). The functional outcomes of APA include modulation of mRNA stability and translation, mRNA localization, and intriguingly, protein localization independent of mRNA localization (2-6). Even though not being as extensively studied as gene transcription, APA is a common phenomenon with at least 70% of mammalian mRNA-coding genes showing APA isoforms (7,8), suggesting that it is one of the major regulatory mechanisms of cellular and molecular dynamics with important functional consequences (1-6). In fact, dysregulation of APA has been associated with disease phenotypes (9) including cancer (10-13), immunological and neurological diseases (14-16).

In recent years experimental methods and computational tools for detecting APA have developed rapidly, allowing the detection of transcriptome-wide APA events on a scale that was not possible previously (9,13). Of interest, there are now computational algorithms that enable detection of APA events directly from standard poly(A)-enriched RNA-seq data, allowing the generation of novel hypotheses and observations using existing RNA-seq datasets from large databases. For example, using DaPars, an algorithm that detects *de novo* PAS proximal to the last exon by modeling read counts of RNA-seq, investigators analyzed data from The Cancer Genome Atlas Project (17) and observed a global shortening of 3’UTRs in tumors compared to normal tissue (11,12). Similar approach was also applied to analyze RNA-seq datasets generated by the Genotype-Tissue Expression (GTEx) project (18) for identification of global APA across different types of human tissues (19-21). Most interestingly, by integrating genome-wide single-nucleotide polymorphism (SNP) genotyping data with APA, investigators were able to discover quantitative trait loci associated with APA (3’aQTLs), many of which had been associated with disease phenotypes but with functions unexplained by expression QTLs (eQTLs) (20,21). Several other biological processes have been reported to affect global APA including cell proliferation (22), cell differentiation (23), and viral infection (24). However, the effects on APA of hormones and their drug analogs, which have mainly been studied for their effects on gene transcription and expression, remain largely unknown with the exception of a case report for estrogens (25). In the present study, we systematically characterized transcriptome-wide the role of glucocorticoids, a major class of hormones, in the regulation of APA.

Clinically, glucocorticoids represent a class of medication used to treat a variety of inflammatory and autoimmune diseases (26,27). While the role of the glucocorticoids in activating the glucocorticoids receptor (GR) to become a transcription factor has been well-characterized, to our knowledge there have been no studies of their regulation of APA. To characterize transcriptome-wide glucocorticoid-dependent APA, we analyzed 90 standard RNA-seq datasets that we had generated from 30 human B-lymphoblastoid cell lines (LCLs) before and after glucocorticoid treatment (28). We discovered a group of immune-specific genes with glucocorticoid-mediated APA that appeared to be functionally distinct from those with glucocorticoid-mediated alterations in expression. We observed that one possible mechanism underlying glucocorticoid-mediated APA involved GR-dependent regulation of RNA-binding proteins (RBP), an important group of APA regulators (29). We also uncovered non-coding genetic risk sequence variants that potentially behaved as alternative polyadenylation quantitative trait loci (3’aQTL), but only after glucocorticoid exposure, some of which had previously been associated with glucocorticoid-related disease phenotypes during genome/phenome-wide association studies.

## MATERIALS & METHODS

### Generation of a 300 LCL genomic panel

Three hundred lymphoblastoid cell lines (LCLs) were obtained from the Coriell Institute and were genotyped with the Illumina HumanHap550K and HumanExon510S-Duo Bead Chips in our laboratory, information that was combined with data generated at the Coriell Institute using the Affymetrix Human SNP Array 6.0. Those genotype data were deposited in the National Center for Biotechnology Information Gene Expression Omnibus under accession number GSE23120 (30).

### Identification of APA events from RNA-seq data

Thirty LCLs with similar levels of GR expression selected from the 300-LCL panel were exposed to cortisol, an endogenous GR ligand, followed by the addition of CORT108297 (C297), a selective GR modulator that could antagonize cortisol activity for 9 hours. RNA-seq for these samples were obtained as described previously (28). The RNA seq data were deposited in the National Center for Biotechnology Information Gene Expression Omnibus under accession number GSE185941. RNA-seq fastq files obtained for the A549 lung carcinoma epithelial cell line after 12 hours of dexamethasone treatment were downloaded from the ENCODE portal (series numbers ENCSR632DQP for control and ENCSR154TDP for treatment). A total of four replicates for each treatment were also downloaded. The fastq files were mapped to the human genome GRCh38 (hg38) using STAR (31). Bed files from each RNA-seq dataset were generated using STAR (31). Identification of APA events was performed with DaPars v2.0 (21) using cutoff values for coverage of 10 and a fit value of 10 using the GENECODE v38 human genome gene annotation. Changes of percentage for the distal poly(A) site usage index (PDUI) between drug and vehicle treatments were conducted in R using a two-sided Wilcoxon matched pair signed rank test. Significance was defined as: (1) FDR less than 0.05 across the 30 LCLs and (2) the fact that the change was reversed by the antagonist C297.

### Identification of differentially expressed genes (DEGs) from RNA-seq

Raw counts were generated from the RNA-seq bam files with the Python package “HTseq” (32) and differential expression analysis was conducted with the R package “EdgeR” (33). Only genes that passed CPM of 32 were included in the analysis. Pathway analysis was conducted with EnrichR (34) both for genes with glucocorticoid-mediated APA and genes with glucocorticoid-mediated expression. Available bed files for cortisol-regulated RBPs were then downloaded from the Encyclopedia of DNA Elements (ENCODE) Project (39). The binding sites of these RBPs were overlapped with 3’UTR data containing the cortisol-mediated APA sites for each gene using bedtools (35).

In addition, precise transcriptional regulation of DEGs in the 30 cells was interrogated using previously generated epigenomic datasets including GR-targeted ChIP-seq obtained in the presence or absence of glucocorticoids, HiChIP targeting H3K27ac, a histone mark associated with active enhancers and promoters, generated with or without exposure to glucocoricoids, and a 25-chromatin state prediction model (28). To identify RBPs that were directly regulated by GR through GR-mediated enhancers, cortisol-dependent RBPs with loops connecting them to cortisol-dependent ChIP-seq peaks were identified using the R packages “GenomicRanges” (36) and bedtools (35). The chromatin states of the cortisol-dependent ChIP-seq peaks were also identified based on the 25-chromatin state prediction model. The sequencing data for these datasets were deposited in the National Center for Biotechnology Information Gene Expression Omnibus under accession number GSE185941 (28).

### Cell culture and drug treatment during validation experiments for glucocorticoid-mediated APA observed for the gene *LY6E*

LCL GM17268 were cultured in RPMI 1640 supplemented with 15% FBS and 1% penicillin/streptomycin. Prior to glucocorticoid treatment experiments, all cells were grown in 5% charcoal-stripped media for 48 hours. The cells were then treated with: (1) vehicle (dimethyl sulfoxide (DMSO) 0.1% and ethanol 0.1%), (2) hydrocortisone 100nM (Sigma Aldrich, dissolved in ethanol) plus DMSO 0.1%; or (1) vehicle, and 100nM dexamethasone (Sigma, water soluble). Treatment time was optimized to achieve the greatest observed effect of glucocorticoid on APA at 3hrs, 6hrs, 9hrs, and 12hrs (data not shown). Since cortisol-induced APA changes were most striking at 9 hours, we repeated the experiments with different dosages of the drugs, specifically, 0nM, 1nM, 10nM, 100nM, and 1000nM after 9 hours of incubation.

### cDNA synthesis

The cells were pelleted and harvested for RNA extraction. Total RNA was extracted with the Quick-RNA Miniprep kit (Zymo, Cat# R1055) per the manufacturer’s instructions. A total of 2μg of total RNA was used for cDNA synthesis using the SuperScript™ III First-Strand Synthesis System (Thermo Fisher, Cat# 18080-093) and poly(A) oligo(dT)_20_ (Thermo Fisher, Cat# 18418020) per manufacturer’s instruction. Briefly, a mixture of 1μL of oligo(dT)_20_ (50 μM), 1μL of 10mM dNTP mix (Thermo Fisher, Cat# 18427088), 2μg of total RNA, and distilled water to 13μL was heated to 65°C for 5 minutes and incubated on ice for 1 minute. This step was followed by the addition of 4μL 5X First-Strand Buffer, 1μL 0.1M DTT, 1μL RNAse Inhibitor (Cat # 10777-019, 40 units/ μL), and 1μL of SuperScript™ III reverse transcriptase (200 units/μL). This mixture was then incubated at 25°C for 5 minutes, 50°C for 45 minutes, and 70°C for 15 minutes. For removal of RNA complementary to the cDNA, 1μL of RNase H (Ambion, Cat # AM2293) was incubated with the mixture at 37°C for 20 minutes. The cDNA was then ready for the PCR step.

### Quantitative Reverse Transcription Polymerase Chain Reaction (qRT-PCR)

Primer sequences targeting the predicted APA site in *LY6E* were designed using Primer3 software. The following two primers were used to perform this experiment: LY6E_3UTR_1 Forward: ACAGCCTGAGCAAGACCTGT, Reverse CGCACTGAAATTGCACAGAA, and LY6E_3UTR_2 Forward FAATGTTGGTGTGGCTTCCAT, Reverse CAGCAGGCTCAGCAGCAG. The product from the cDNA synthesis experiment was diluted 200 times before qRT-PCR, and the primers were diluted to a final concentration of 0.4 μM. Reaction for qRT-PCR was performed using Power SYBR™ Green PCR Master Mix (Applied Biosystems Inc., Cat# 43-676-59). Analysis of qRT-PCR data was conducted using the 2^*-*ΔΔCT^ method (37).

### Analysis of glucocorticoid-mediated alternative polyadenylation quantitative trait loci

1.3 million genotyped single nucleotide polymorphisms (SNPs) were filtered based on Hardy-Weinberg Equilibrium (P > 0.001) with genotyping call rates of more than 95%. All SNPs which had only 1 or 2 cell lines with homozygous variant genotypes among the 30 cell lines studied were excluded from further study. PDUI for all treatment conditions were normalized to vehicle by genotypes, thus cancelling out genotype effects at baseline. APA events without adequate coverage in more than one-third of the 30 LCLs and with no change in more than half of the 30 LCLs were excluded. Analysis of quantitative trait loci for delta PDUI was conducted with the R package “Matrix eQTL” (38) using ANOVA model. A total of 515,177,592 associations were conducted, and *P*-values with significant trends were defined as 5.16 × 10^−8^. Those SNPs were then overlapped with SNPs documented in the GWAS Catalog (https://www.ebi.ac.uk/gwas/) and with UKBiobank data (http://pheweb.sph.umich.edu/) for potentially significant clinical associations. Significant *P*-value cutoffs were defined by the databases queried.

## RESULTS

### Glucocorticoids mediated global APA in human LCLs

Thirty LCLs of differing genomic backgrounds were treated with vehicle, 100 nM cortisol (a GR agonist) and 100 nM cortisol plus 100 nM of CORT108297 (C297, a GR antagonist). Treatment conditions such as dose and incubation time were optimized before sequencing, and the treatment effect was confirmed for these datasets by expression quantification of prototypic GR-targeted genes such as *FKBP5* and *TSC22D3*. Specifically, the mRNA levels for both of these genes were significantly induced after cortisol treatment and those inductions returned to baseline after C297 was added to the treatment (28). We then analyzed these 90 RNA-seq datasets to identify transcriptome-wide APA events using the computational algorithm DaPars v.2.0 (21). DaPars gave an output of the percentage of distal poly(A) site usage index (PDUI) for each gene transcript, together with the position of the predicted proximal polyadenylation site (PAS). A higher value of PDUI corresponded to a higher percentage of transcripts with longer 3’UTRs or more distal PAS. After PDUI values were obtained for each gene in each sample, a total of 696 genes that displayed APA had been detected. Of these 696 genes, we found that cortisol regulated APA dynamics for 54 genes (**Figure 1A, Supplementary Table 1**) (two-sided Wilcoxon matched pair signed rank test; FDR < 0.05 across 30 LCLs). Importantly, those cortisol-mediated changes in APA were reversed after C297, ie, after antagonist treatment, confirming that the observed APA changes were dependent on glucocorticoid treatment. For example, cortisol decreased the percentage of distal PAS usage by 30% for LY6E mRNA, a gene that encodes lymphocyte antigen 6 family member E, resulting in a shortening of the LY6E 3’UTR (**Figure 1B-C)**. The addition of C297 reversed this cortisol-mediated repression back to baseline (**Figure 1B-C**). Interestingly, we found that cortisol treatment did not change LY6E mRNA levels in our LCLs but that it did mediate 3’UTR APA to reduce the percentage of transcripts with long 3’ UTRs (**Figure 1B**). To validate the glucocorticoid-mediated APA observations for *LY6E*, we quantified the long LY6E 3’UTR transcripts by qRT-PCR using two different sets of primers targeting the LY6E 3’UTR region that covered the proximal PAS. We observed a dose-dependent repression by cortisol, and we replicated that dose-dependent regulation with dexamethasone, a potent synthetic glucocorticoid that is often used in the clinic (**Figure 1D**).

**Figure 1.**
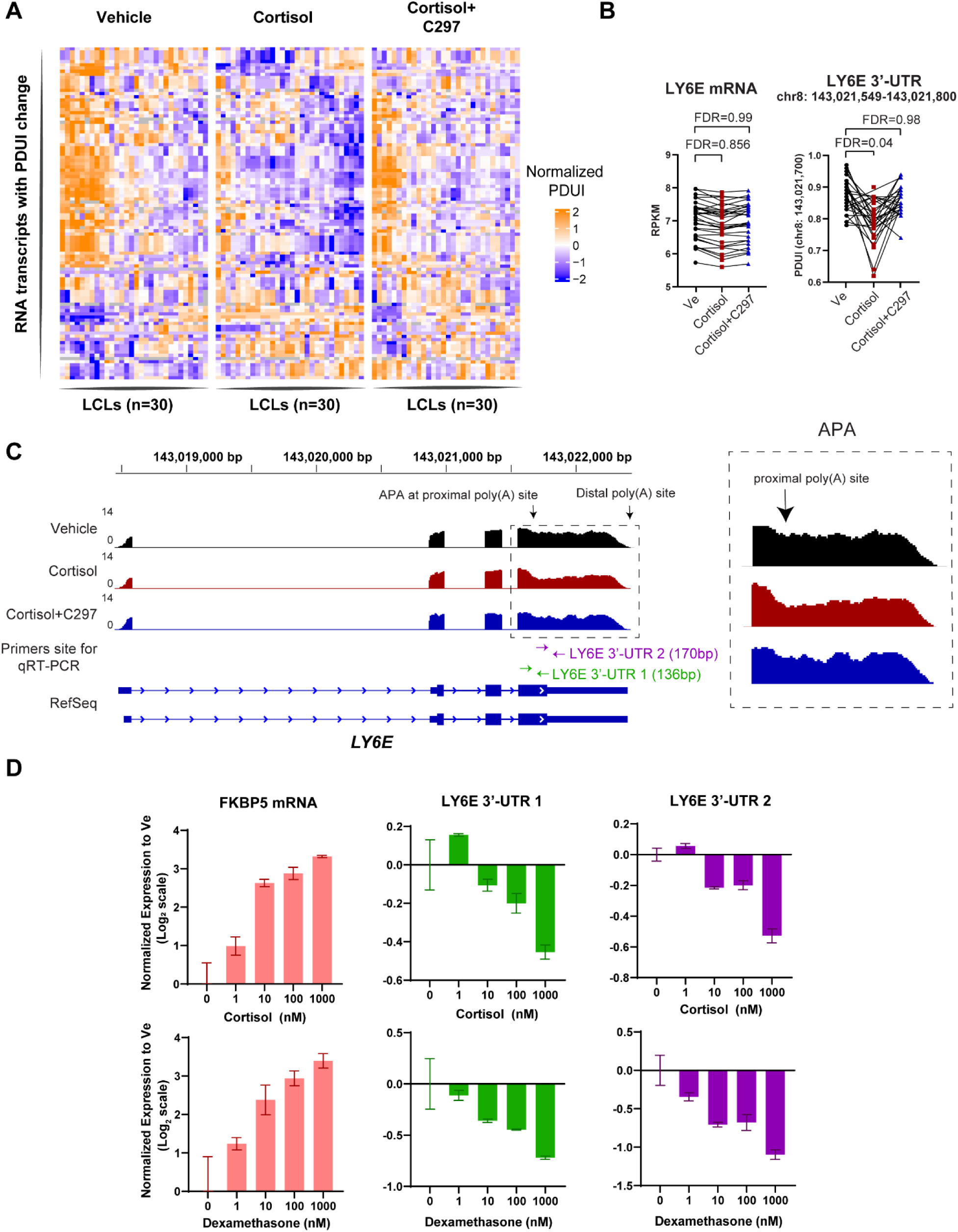
Glucocorticoids regulate transcriptome-wide APA in human LCLs. (**A**) Heatmap showing changes in the Percentage of Distal APA Usage (PDUI) across treatment conditions. Cortisol is an agonist and C297 is an antagonist. Each row represents a cell line, and each column represents PDUI at a particular APA site. (**B**) *LY6E* mRNA expression level and APA change as measured by PDUI after glucocorticoid treatment from RNA-seq data across 30 LCLs. Each dot represents a cell line, with lines connecting the cellular status at each treatment point. (**C**) An Integrative Genomic Viewer (IGV) plot of the cortisol-mediated LY6E 3’-UTR APA site. Each track represents a bedgraph file for RNA-seq reads. (**D**) Dose-dependent effect of 2 different glucocorticoids on the APA site from (**B-C**) as measured by qRT-PCR using two different primers.

### Glucocorticoids mediated APA through transcriptional regulation of RNA-binding protein in human LCLs

When we conducted pathway analysis of 1362 cortisol-responsive genes from our RNA-seq results (FDR < 0.05), we found that “RNA-binding protein” (RBP) was the most enriched pathway (adjusted *p*-value = 7.39E-04), raising the possibility that cortisol might regulate downstream RBPs with APA-regulating properties (**Figure 2A**). Therefore, we hypothesized that glucocorticoid-activated GR might regulate the expression of RBPs with known APA-regulating properties, which in turn might drive the changes observed for APA. From the pathway analysis, we observed that 142 proteins with RNA-binding properties were regulated by cortisol and that their cortisol-mediated expression was reversed by C297 (**Figure 2B**). To identify which of those 142 DEGs were directly regulated by GR, we consulted GR-targeted ChIP-seq and H3K27ac HiChIP datasets that we had previously generated in LCLs both with and without cortisol treatment, together with a 25-chromatin-state prediction model (28). We found that 27 (18%) of the 142 glucocorticoid-regulated RBPs had H3K27ac HiChIP loops that connected them with one or more cortisol-induced GR peaks, suggesting direct regulation through cortisol-mediated enhancers (**Figure 2C)**.

**Figure 2.**
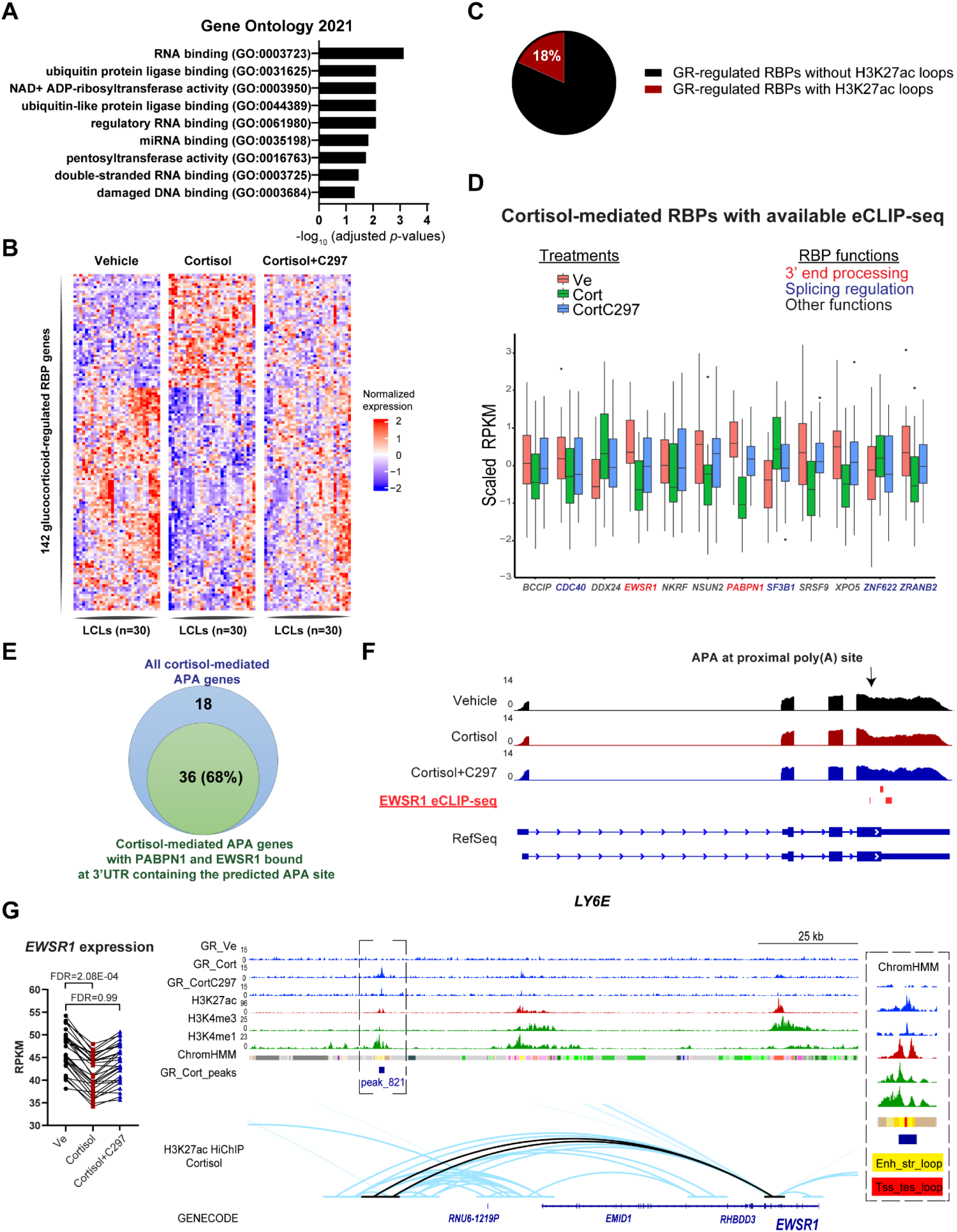
Potential mechanisms underlying glucocorticoid-mediated APA in human LCLs. (**A**) Pathway analysis of all cortisol-responsive genes in the 30 LCLs studied revealed that RNA-binding was the most enriched pathway based on the Gene Ontology 2021 database. (**B**) Heatmap depicting changes in expression of 142 RBPs from the top pathway in (**A**) across treatment conditions. Each column represents a cell line, and each row represents expression of a gene. (**C**) Pie chart depicting the percentage of genes encoding RBPs from **(B**) and **(C**) that are regulated directly by GR through GR-mediated enhancers that loop to genes to transcriptionally regulate their expression. (**D**) Twelve cortisol-mediated RBPs that have genome-wide RNA-binding patterns available on ENCORE through eCLIP assay. The Y axis represents z-scores for gene expression after exposure to vehicle, CORT or CORT/C297. (**E**) The number of genes with cortisol-mediated APA that have RBPs with known APA-regulating properties that bind at their 3’-UTRs containing the predicted APA site based on data from (**D**). (**F**) An IGV plot showing the *LY6E* 3’UTR, a glucocorticoid-mediated APA example from **Figure 1**, in which an RBP with known APA-regulatory properties, *EWSR1*, binds to multiple regions surrounding the predicted APA site. (**G)** *EWSR1* expression is repressed by cortisol via a distal enhancer, as shown in the IGV plot. Loops that directly interact between the GR-binding site and the *EWSR1* promoter are colored black. Chromatin state abbreviations: Enh_str_loop: Enhancer with strong looping properties; Tss_tes_loop: POLR2 initiation & stop with enhancer-looping property.

We then explored the function of the 142 cortisol-regulated RBPs by consulting enhanced crosslinking and immunoprecipitation sequencing (eCLIP-seq) datasets for more than 70 RBPs and their annotated functions that had been generated by the ENCODE project (39). We found that 12 of those cortisol-regulated RBPs had information for transcriptome-wide RNA-binding sites readily available in two human cell lines (**Figure 2D;** see **Supplementary Table 2** for specific ENCODE series numbers). Two of those 12 RBPs, EWSR1 and PABPN1, are known to bind to 3’UTRs and to regulate APA (40,41). Specifically, PABPN1 is known to suppress APA or prevent 3’UTR shortening (40), while EWSR1 is part of the FET RNA binding proteins family that can regulate APA depending on their binding sites (41). The other 10 cortisol-regulated RBPs such as CDC40, ZRANB2, DDX24, NKRF, SF3B1 and NSUN2 are known to play diverse roles in RNA processing (42-47), but their roles—if any—in the regulation of APAs have yet to be determined.

When we integrated the eCLIP-seq data for APA-regulating RBPs PABPN1 and EWSR1, as described above, with our APA analysis, we found that 68% of 54 genes with GR-mediated APA events were bound by either EWSR1 or PABPN1 at 3’UTRs containing the alternative PAS (**Figure 2E)**. For example, based on eCLIP-seq data, EWSR1 bound to multiple sites in the *LY6E* 3’-UTR surrounding the predicted proximal PAS (**Figure 2F)**. EWSR1 was downregulated by cortisol via a cortisol-induced GR enhancer that loops across 100kb to the *EWSR1* promoter (**Figure 2G**), possibly causing the observed repression of the distal PAS usage. Taken together, these observations supported our hypothesis that glucocorticoids can mediate APA, at least in part, through the transcriptional regulation of RBPs with APA-regulating properties.

GR is known to act as an RBP itself (48). However, a previous study reported that, in general, GR binds to the 5’UTR rather than the 3’UTR of mRNAs and that GR-RNA binding is independent of AU-rich elements, an RNA-binding motif that is often found in 3’UTRs (48). This observation suggests that the RNA-binding property of GR might have limited effect on the glucocorticoid-dependent APA that we observed.

### Genes differentially expressed and genes alternatively polyadenylated by glucocorticoids are functionally distinct

As described above, we identified 54 glucocorticoid-mediated APA genes, a much smaller number than the 1,362 glucocorticoid-mediated DEGs in these same LCLs. Furthermore, we noticed that only 5 of those 54 APA genes overlapped with the 1,362 DEGs (**Figure 3A**). Therefore, we asked what the function(s) of the glucocorticoid-mediated APA genes might be and how they might differ from those of the glucocorticoid-mediated DEGs. It is well known that one of the mechanisms underlying the immunosuppressant actions of glucocorticoids is medicated through the inhibition of interleukin and cytokine signaling pathways (49). For example, glucocorticoids induce the expression of glucocorticoid-induced leucine zipper (*GILZ*, or *TSC22D3*), an inhibitor of NF-kB (50) and they inhibit the expression of pro-inflammatory cytokines and chemokines (49). We observed similar effects of cortisol in our LCLs, as demonstrated by pathway analysis of cortisol mediated DEGs (**Figure 3B, Supplementary Table 3**). However, we found that genes which displayed glucocorticoid-mediated APA were primarily enriched in viral translation-related pathways, with *p-*values more significant than those for pathways enriched for cortisol-mediated DEGs despite a smaller number of this class of genes (**Figure 3C-D)**. That observation was consistent with previous studies which demonstrated that APA can play a crucial role in antiviral immune response (24,51). Indeed, *LY6E*, a gene showcased in previous paragraphs, represented an example of a glucocorticoid-mediated APA gene with emerging roles in a wide spectrum of viral diseases. Specifically, *LY6E* encodes a glycosyl-phosphatidyl-inositol-anchored cell surface protein that plays an important role in immunological regulation including modulation of viral infection by coronaviruses such as SARS-CoV-2 (52-55). Specifically, LY6E has been shown to be an antiviral immune effector by interfering with spike protein-mediated membrane fusion, protecting a variety of cell types including primary B cells from coronaviruses infection (53,54). The functional consequences for viral biology of the shortening of LY6E 3’UTR after cortisol treatment, however, remains to be determined.

**Figure 3.**
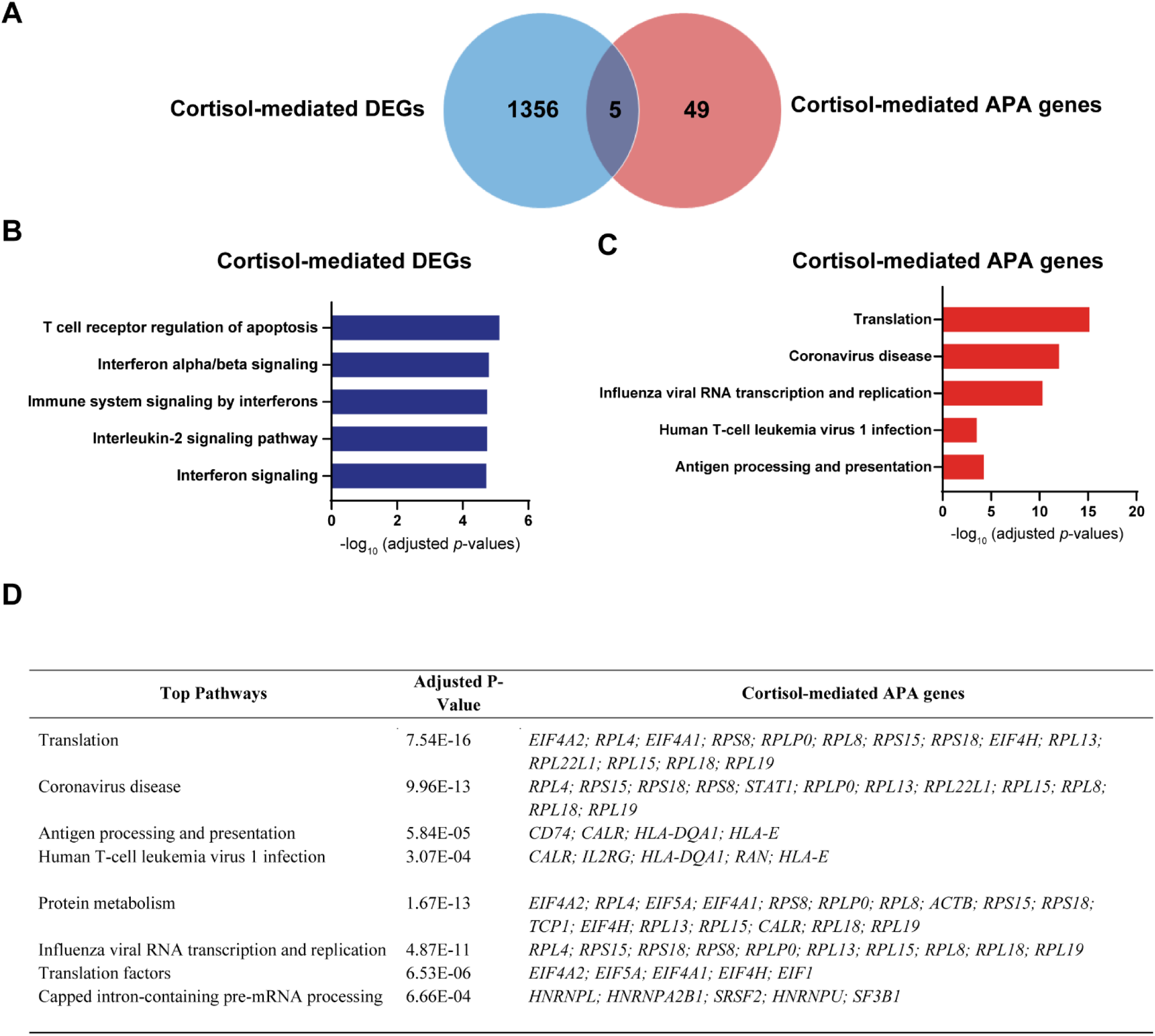
Glucocorticoid-mediated APA is functionally distinct from glucocorticoid-mediated gene expression. (**A**) Overlap of genes with cortisol-mediated APA and cortisol-mediated DEGs in LCLs. (**B-C**) Selected top pathways for genes with cortisol-mediated APA and DEGs in LCLs. (**D**) Specific cortisol-mediated APA genes enriched for in pathways depicted in **(B)**.

### Glucocorticoid-mediated APA is cell-type specific

To determine whether the glucocorticoid-mediated APA effects that we observed in LCLs might apply to other types of cells, we analyzed published RNA-seq datasets from the ENCODE portal generated for A549 lung carcinoma epithelial cells before and after 100nM dexamethasone treatment for 12 hours, a treatment condition comparable to that used in our LCL study (56). Using similar analytical approaches to those which we applied for LCLs, we identified 2448 DEGs after dexamethasone treatment of A549 cells (FDR < 0.05), with 243 genes overlapping with DEGs that we had identified in LCLs (see **Figure 4A**). Interestingly, the most significant pathway enriched for glucocorticoid-regulated DEGs in A549 cells was the DNA-binding pathway (adjusted *p*-value = 3.022e-8) (**Figure 4B**), in contrast to the RNA-binding pathways observed in LCLs, as described above (**Figure 2A**). Using the DaPars v2.0 algorithm, we were able to detect 460 genes with APA events at baseline in A549 cells, genes which overlapped with 56% of the genes with APA that we had detected in LCLs (see **Figure 4C**). However, we did not detect any change in APA events after the dexamethasone treatment of A549 cells (FDR < 0.05) (**Figure 4D)**. Consistent with these observations, much fewer RNA-binding-related genes were differentially expressed in A549 cells than that in LCLs (21 versus 142 genes) after glucocorticoids treatment (**Figure 4A**). Importantly, two genes with known APA-regulating properties and explaining 68% of glucocorticoid-dependent APA observed in the LCLs, *PABPN1* and *EWSR1* (**Figure 2E**), were not differentially expressed in A549 cells after glucocorticoid treatment (**Figure 4A**). This result might explain why no glucocorticoid-mediated APA events were detected in A549 cells. These observations suggest that GR-mediated APA regulation may be highly cell-type specific and that it may occur more often in association with immune-related functions of GR.

**Figure 4.**
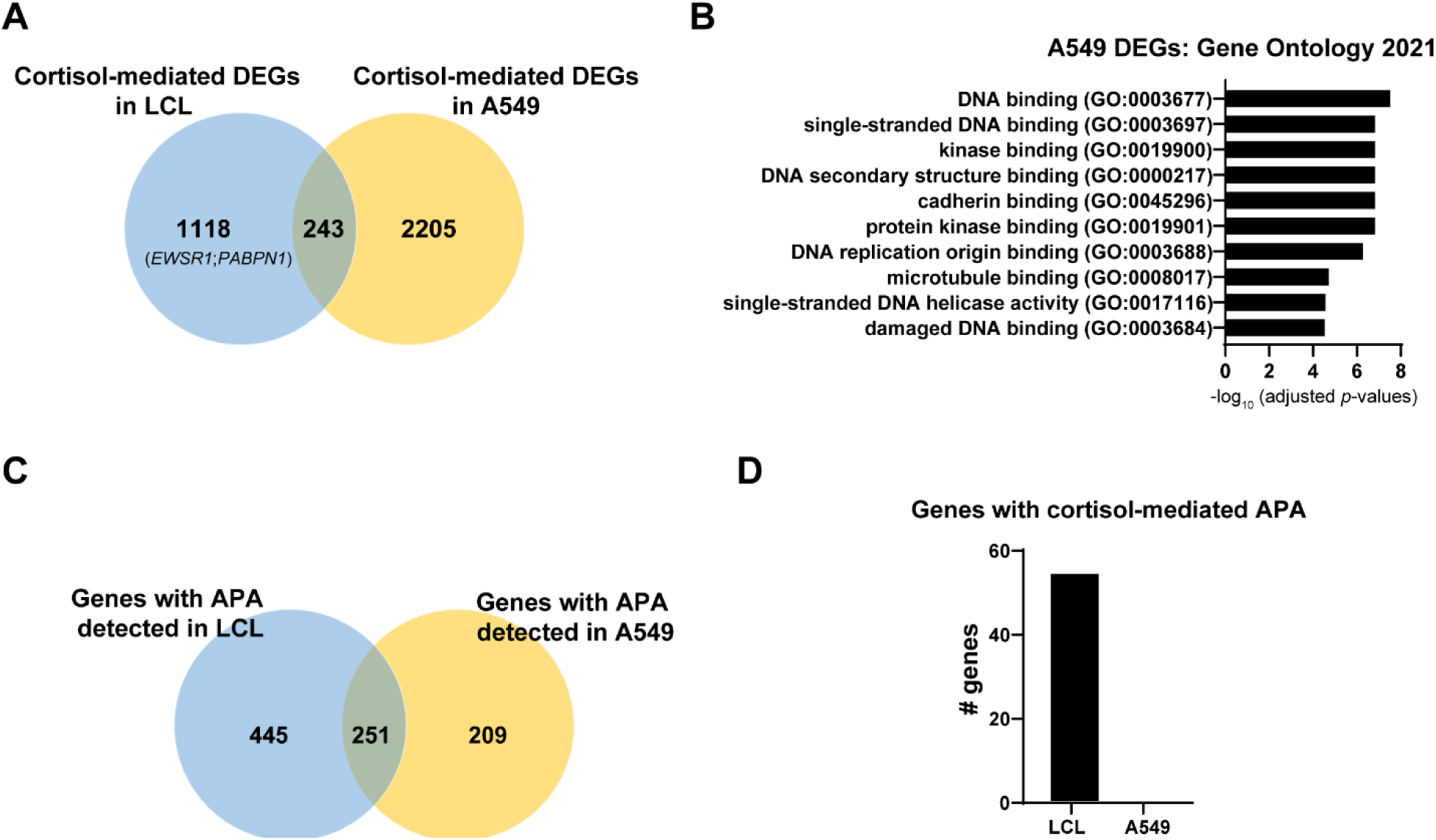
Glucocorticoid-mediated APA is cell-type specific. (**A**) Overlap of differentially expressed genes induced by glucocorticoids in LCLs and in A549 cells. (**B**) Top pathways enriched for DEGs in A549 cells. (**C**) Overlap of genes with APA events detected in LCLs and in A549 cells. (**D**) Number of genes with glucocorticoid-mediated APA in A549 cells and in LCLs.

### Glucocorticoid mediated APA in a genotype-dependent manner: association with clinical phenotypes

Taking advantage of the genome-wide genotype data that we had already generated for our 30 LCLs, we asked whether single-nucleotide polymorphisms (SNPs) might contribute to the observed variation in glucocorticoid-mediated APA. The fact that APA regulation at baseline can be genotype-dependent had already been demonstrated by other investigators based on the concept of 3′UTR alternative polyadenylation quantitative trait loci (3′aQTLs) (21,57). Utilizing the GTEx database, Li et al, 2021 (21) had observed that 3’aQTLs could potentially explain approximately 16.1% of currently known trait/disease-associated genetic variants. Acknowledging that 30 LCLs provided us with limited power, we undertook an exploratory search to identify evidence that SNP loci could be associated with glucocorticoid-mediated APA events, referred to hereafter as glucocorticoid-mediated pharmacogenomic-3’aQTLs (PGx-3’aQTLs). We then explored the possible link of those PGx-3’aQTLs to genetic variants that had been associated with clinical phenotypes. With the 30 LCLs used in this study, we were able to re-capture a series of previously identified 3’aQTL SNPs, such as SNPs in tight linkage disequilibrium (*r*^2^ >0.70) with the rs10954213, a SNP that was associated with APA for the *IRF5* 3’UTR (16,21) (*P* = 4.69 × 10^−8^) (**Figure 5A**).

**Figure 5.**
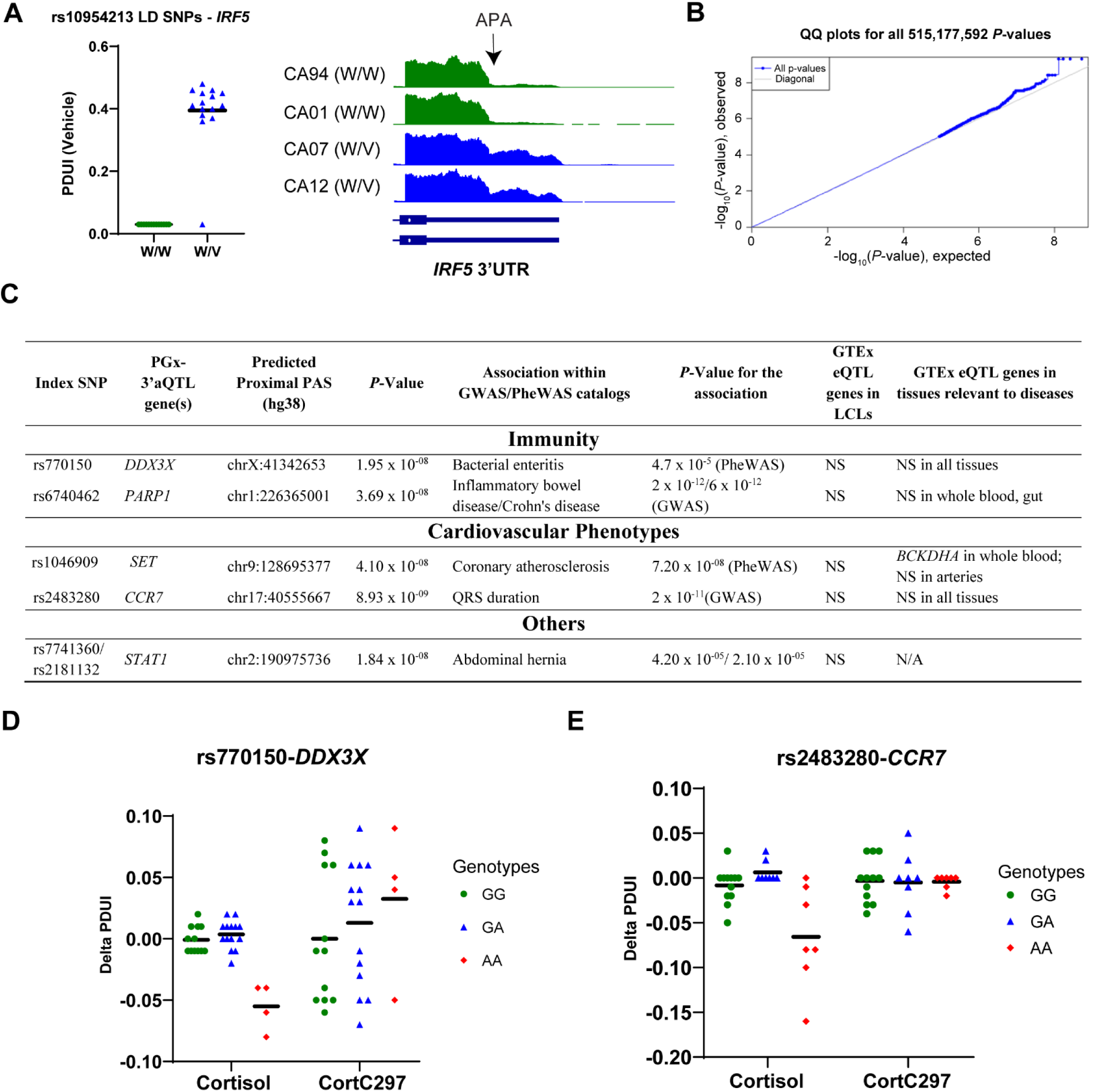
Glucocorticoid mediated APA in a genotype-dependent manner: associations with diseases involving glucocorticoid-signaling. **(A)** The 30 LCLs used in this study captured a previously studied 3’aQTL related to the 3’UTR of *IRF5* through SNPs in the LD block. (**B**) QQ-plots of all SNP-APA gene pairs identified during PGx-3’aQTL analysis indicated that no inflation was observed. (**C**) PGx-3’aQTLs that have been associated with diseases based on previous GWAS/PheWAS results. (**D-E**) Selected examples of PGx-3’aQTL with clinical implications from (**C)** showing allele-dependent and drug-dependent regulation of APA in specific genes.

To identify possible glucocorticoid-mediated PGx-3’aQTL associations, we associated SNP genotypes with changes in percentage of distal APA usage between glucocorticoid treatment and vehicle exposure across our 30 LCLs (see **Methods**). That step identified a total of 66 cortisol-dependent candidate PGx-3’aQTLs (*P* < 5.16 × 10^−8^), all of which no longer exhibited SNP-APA change associations after the addition of the GR antagonist, C297 (*P* ≥ 0.05), demonstrating a cortisol-dependent effect (**Figure 5B, Supplementary Table 4**). To investigate whether these glucocorticoid-dependent PGx-3’aQTLs could potentially explain the function of SNPs previously associated with clinical phenotypes but with unclear functional mechanisms, we overlapped PGx-3’aQTL SNPs with those already reported to be genome-wide significant signals in the GWAS Catalog (58) and/or the UK Biobank PheWAS Catalog (59). We found that five of our PGx-3’aQTLs had been associated with disease phenotypes (**Figure 5C**), four of which were related to known effects of glucocorticoid usage such as bacterial infections (60), inflammatory bowel disease (61,62), and cardiovascular disease (63,64). Furthermore, the function of the genes modulated by the PGx-3’aQTL appeared to be related to these associated phenotypes. For example, rs770150 was associated with bacterial enteritis based on PheWAS (*P* = 4.7 × 10^−05^), and it was found in our study to be a PGx-3’aQTL for *DDX3X*. Specifically, cortisol decreased PDUI in subjects with the AA genotypes but not subjects with other genotypes at this locus, and this repression was reversed by C297 (**Figure 5D**). *DDX3X* encodes a RBP that is known to play a critical role in antimicrobial innate immunity (65,66). Another example involved rs2483280, a SNP previously associated by GWAS with electrocardiographic QRS duration (2.0 × 10^−11^). That SNP was associated with repressed PDUI for *CCR7* in a genotype- and drug-dependent manner (**Figure 5E)**. *CCR7* encodes a chemokine receptor that has been shown to play a role in cardiac function (67,68). While these associations with disease through possible glucocorticoid-dependent mechanisms are intriguing, these SNPs did not disrupt GR binding sites and appeared to act through *trans* mechanisms that remain to be determined and validated in future studies with larger sample sizes.

## DISCUSSION

Our study has provided a series of observations with regard to the impact of glucocorticoids on alternative polyadenylation which, to our knowledge, had not been reported previously. In the clinic, glucocorticoids are used to treat a variety of inflammatory and autoimmune diseases (26,27). Traditional understanding of glucocorticoid mechanisms in immune regulation has been based on their activation of the glucocorticoid receptor (GR), which is then translocated to the nucleus where it binds to DNA to initiate gene transcription (17,18). We demonstrated that glucocorticoids could also regulate global APA for immune-related genes in human LCLs (**Figure 1**), genes that were functionally distinct from those influenced by glucocorticoids on expression level (**Figure 3**). Even though the number of glucocorticoid-dependent APA genes was smaller than that of glucocorticoid-dependent DEGs, glucocorticoid-dependent APA genes were highly enrich in the “viral infection-related” pathways (**Figure 3C**), indicating a more specific function of glucocorticoid-dependent APA. Characterization of glucocorticoid-dependent APA in viral infection might help to better understand the mechanism of glucocorticoids’ therapeutic effects in diseases such as COVID-19 (69,70).

Glucocorticoids were known to play important functional roles in a variety of multiple physiological processes and diseases including autoimmunity (49), osteoporosis (71), mood disorders (72,73), cancer (74), and metabolism (75). However, the therapeutic usage of glucocorticoids remains largely dependent on their function in immunosuppression. Of interest, glucocorticoid-dependent APA, which possibly occurred through glucocorticoid-mediated transcriptional regulation of RBPs with APA-regulating properties (**Figure 2**), appears to be cell-type specific (**Figure 4**). Although comparable numbers of APA genes were identified in both LCLs and A549 lung cancer cells (**Figure 4C**), none of the glucocorticoid-dependent APA genes were identified in the A549 cell line **(Figure 4D**), raising the possibility that glucocorticoid-dependent APA could be more often immune-related. This possibility requires further investigation with more cell lines and tissue types, which, once confirmed, might expand our understanding of the mechanism(s) of glucocorticoids’ immunosuppressive properties. However, the present study did not determine the functional consequences of these APA changes in terms of molecular and cellular dynamic changes induced by glucocorticoids. While mRNA stability is one of the known functional outcomes of APA (1), we found that glucocorticoid-mediated changes in APA were only associated with changes in gene expression in a small minority of DEGs in LCLs.

We also observed trends that glucocorticoids could regulate APA in a genotype-dependent manner, a phenomenon that we referred to as “PGx-3’aQTL” (**Figure 5**). PGx-3’aQTLs could represent a novel type of context-dependent SNPs for which pharmacological or physiological reagents can “unmask” the function of silent non-coding SNPs with previously unknown functional roles. They could also potentially explain an aspect of gene × environment interaction in disease risk and variation in drug response that otherwise could not be explained by eQTLs or context-dependent eQTLs. Unlike diseases associated with glucocorticoid-mediated PGx-eQTLs which spanned a broad spectrum of glucocorticoid-related diseases as identified in our previous study (28), diseases or pathological states associated with glucocorticoid-mediated PGx-3’aQTLs appeared to be limited to or focused on immune-related phenotypes—at least in LCLs. This observation is consistent with the fact that glucocorticoids regulate APA in a cell-type specific manner and appeared—in LCLs—to do so primarily for genes with immune-related functions. While these observations are intriguing, they need to be validated with larger sample size.

In conclusion, our study has revealed a series of novel aspects of genomic regulation by glucocorticoids, particularly their impact on RNA polyadenylation with its possible role in disease risk, aspects that should be explored for other hormones and nuclear receptors beyond GR.

## Data availability

The genotyping data for this study were deposited in Gene Expression Omnibus under accession number GSE23120. All other sequencing data were deposited under accession number GSE185941.

## Supplementary Data

**Supplementary Table 1 (Excel Sheet)**. Genes with APA changes after glucocorticoids treatment in 30 human LCLs

**Supplementary Table 2 (Excel Sheet)**. ENCODE series number of eCLIP datasets for 12 RBPs used in this study.

**Supplementary Table 3**. Cortisol-mediated DEGs enriched in pathways extracted from databases KEGG 2021 and Bioplanet 2019 from EnrichR server.

**Supplementary Table 4 (Excel Sheet)**. Cortisol-mediated PGx-3’aQTLs in 30 human LCLs.

## Author contributions

Conceptualization: DL, TTLN, RMW; Investigation: TTLN, DL, LZ; Data Curation: HG, LWei; Formal Analysis: TTLN, HG, ZY; Methodology: JHL, TO; Resources: LWang, TO; Supervision: RMW, LWang, TO; Funding Acquisition: RMW and LWang; Visualization: TTLN; Data Interpretation: TTLN, DL, HG, JY, LWang, TO, RMW; Writing – original draft: TTLN, DL, RMW; Writing – review & editing: JHL, TO.

## Acknowledgements

We thank the ENCODE Consortium for their generation of the epigenomic datasets used in this study.

