## Supplementary table 3 for "Glucocorticoids Mediate Transcriptome-wide Alternative Polyadenylation: Mechanisms and Functional Implications"

**Supplementary Table 1 (Excel Sheet). Genes with APA changes after glucocorticoids treatment in 30 human LCLs**

**Supplementary Table 2 (Excel Sheet). ENCODE series number of eCLIP datasets for 12 RBPs used in this study.**

**Supplementary Table 3**. Cortisol-mediated DEGs enriched in pathways extracted from databases KEGG 2021 and Bioplanet 2019 from EnrichR server.

| Top Pathways | Adjusted P-Value | Genes with cortisol-mediated expression |
|---|---|---|
| T cell receptor regulation of apoptosis | 7.64E-06 | *CLIC5; CSF3; BTG1; HCCS; ITGAL;* ***NR3C1****; TNF; CCND3; CASP6; MYB; PIM1; RAC1; HRAS; IER2; MAP4K1; DUSP1; TSC22D3; RHOH; DYNLL1; PSMA5; TLR1; CH25H; PSMA6; CLIP1; MMRN1; EWSR1; IRF4; IRF7; LTA; PSME2; TLR7; LAP3; TLR6; LTB; BIRC3; FAIM; MAX; GLO1; FAM35A; COPS7B; HTT; CSF2RB; RASGRP2; SLC5A3; FOXO1; RELB; STRN3; PRDX1; TP53BP2; PMAIP1; BMF; MAPK6; APOL3; S100A10; ZBTB18; MAP3K2; HSPA9; FDPS; SLC16A1; EGR3; STAT1; FEM1B; TREX1; MX1; HUWE1; CFLAR; UBE2A; CLK1; TBL3; SCD; DNAJA3; JMY; ERCC2; CYCS; IL7R; PDE7A; MAP3K14; ADA; TMEM9B; CNIH4* |
| Interferon alpha/beta signaling | 1.59E-05 | *PTPN1; RNASEL; STAT1; MX2; MX1; ADAR; ISG15; IFI35; IFIT1; USP18; ISG20; SOCS1; OAS1; IRF4; OAS2; IRF7; IRF6; GBP2; XAF1* |
| Immune system signaling by interferons, interleukins, prolactin, and growth hormones | 1.81E-05 | *RNASEL; EIF4E3; UBA7; CSF2RB; ADAR; IFI35; NOD2; IFI30; IFIT1; USP18; SOCS1; UBB; TRIM25; GBP2; HRAS; CAMK2G; LYN; PTPN1; MAP2K1; CISH; STAT1; IFNGR2; MX2; MX1; EIF2AK2; ISG15; YWHAZ; PML; NUP93; ISG20; OAS1; PIK3CA; IRF4; OAS2; TNIP2; NUP50; IRF7; PIN1; GRB2; IRF6; IL6ST; IL7R; XAF1; EIF4G3; MYD88* |
| Interleukin-2 signaling pathway | 1.81E-05 | *FCGBP; CSF1;* ***NR3C1****; IFIT1; TNF; TM7SF2; RGS3; RGS1; MYB; PIM1; TNFSF10; BCAP29; FAM53B; ENPP4; PIAS4; SLC38A1; MAP2K1; CISH; HMHA1; PASK; MTHFD1; STIM1; KARS; IL6ST; YTHDC2; TCF7; CSF2RB; RASGRP2; FOXO1; ZFP36L2; TMEM243; SOCS1; RDH11; STK38; PMAIP1; BMF; KCNN4; SLAMF1; LYN; GOT1; FUCA1; IFI44; NMI; EIF2S2; SQLE; FUBP1; ID3; GRB2; IL7R; ADA; BTG1; MAST3; PLEK; PTPN22; CMAHP; ITGB7; HRAS; WARS; KLF13; DUSP1; TSC22D3; BCKDHB; BTN3A3; PARP8; PIK3CA; IRF4; LTA; GARS; TOP1; TLR6; LTB; VAMP5; BIRC3; NFAT5; PCNA; RALB; AHNAK; SRF;* |

| | | |
|---|---|---|
| | | *CXCR4; DENND3; MNT; SAMM50; NFIL3; E2F3; IL12RB1; CD55; POLR2M; STAT1; CD70; TXNRD1; MX1; CFLAR; KLF6; P2RX5; CAMK4; PDCD4; ESYT2; PDE7A; LPIN2* |
| Interferon signaling | 1.92E-05 | *RNASEL; EIF4E3; UBA7; ADAR; IFI35; IFI30; IFIT1; USP18; SOCS1; UBB; TRIM25; GBP2; CAMK2G; PTPN1; STAT1; IFNGR2; MX2; MX1; EIF2AK2; ISG15; PML; NUP93; ISG20; OAS1; IRF4; OAS2; NUP50; IRF7; PIN1; IRF6; XAF1; EIF4G3* |

**Supplementary Table 4 (Excel Sheet). Cortisol-mediated PGx-3'aQTLs in 30 human LCLs.**
